# Studying magnetic susceptibility, microstructural compartmentalisation and chemical exchange in a formalin-fixed ex vivo human brain specimen

**DOI:** 10.1101/2021.07.30.454493

**Authors:** Kwok-Shing Chan, Renaud Hédouin, Jeroen Mollink, Jenni Schulz, Anne-Marie van Cappellen van Walsum, José P. Marques

**Affiliations:** Donders Institute for Brain, Cognition and Behaviour, Radboud University, Nijmegen, The Netherlands; Empenn, INRIA, INSERM, CNRS, Université de Rennes 1, Rennes, France; Department of Medical Imaging, Anatomy, Donders Institute for Brain, Cognition and Behaviour, Radboud University Medical Center, Nijmegen, The Netherlands

**Keywords:** Quantitative susceptibility imaging, phase imaging, ex vivo imaging, microstructure, white matter

## Abstract

**Purpose:** Ex vivo imaging is a preferable method to study the biophysical mechanism of white matter orientation-dependent signal phase evolution. Yet, how formalin fixation, commonly used for tissue preservation, affects the phase measurement is not fully known. We, therefore, study the impacts of formalin fixation on magnetic susceptibility, microstructural compartmentalisation and chemical exchange measurement on human brain tissue.

**Methods:** A formalin-fixed, post-mortem human brain specimen was scanned with multiple orientations with respect to the main magnetic field direction for robust bulk magnetic susceptibility measurement with conventional quantitative susceptibility imaging models. Homogeneous white matter tissues were subsequently excised from the whole-brain specimen and scanned in multiple rotations on an MRI scanner to measure the anisotropic magnetic susceptibility and microstructure-related contributions in the signal phase. Electron microscopy was used to validate the MRI findings.

**Results:** The bulk isotropic magnetic susceptibility of ex vivo whole-brain imaging is comparable to in vivo imaging, with noticeable enhanced non-susceptibility contributions. The excised specimen experiment reveals that anisotropic magnetic susceptibility and compartmentalisation phase effect were considerably reduced in formalin-fixed white matter tissue.

**Conclusions:** Despite formalin-fixed white matter tissue has comparable bulk isotropic magnetic susceptibility to those measured via in vivo imaging, its orientation-dependent components in the signal phase related to the tissue microstructure is substantially weaker, making it less favourable in white matter microstructure studies using phase imaging.

## Introduction

Quantitative susceptibility mapping (QSM) is a physics-driven method to the study magnetic properties of biological tissues (1). Some features that differentiated it from conventional MR relaxometry include the field strength independence of the derived maps, relying on spatial deconvolution and its ability to distinguish paramagnetic and diamagnetic substances since they produce opposite contrasts. QSM is commonly performed on gradient echo phase data owing to its direct relationship to magnetic field variations (2,3).

One major QSM research challenge is to understand the mechanism of phase evolution in white matter (WM) (4,5). In deep grey matter (GM), strong correlations between QSM and iron concentration have been demonstrated (6). Yet, in WM, the abundance of diamagnetic myelin (relative to water) would have suggested a strong QSM contrast relative to cerebrospinal fluid (CSF) (4,5). The lack of this strong contrast has been attributed to various biophysical phenomena (7,8). The lipid-rich myelin bilayer sheath encapsulating the highly-ordered axons in WM results in anisotropic susceptibility (7–12). Additionally, water protons exist in various microstructural environments (13), namely myelin water, intra-axonal water and extra-axonal water, which can have different signal decay rates and frequency shifts depending on the composition of the tissue and the fibre orientation with respect to the main magnetic field (B_0_) (14–17), which also make the signal phase not representing the average magnetic field in a voxel. The chemical exchange of protons between macromolecules and water can also introduce a further frequency shift of the MR signal (18,19). Disentangling the origins of WM phase contrast can improve our understanding of QSM and provide new means to account for their impact in QSM.

As WM phase contrast is orientation-dependent, studying its properties requires data acquired with different orientations to B_0_. Subject compliance limits the range of angles that can be obtained in vivo (unnatural posture inside the scanner). Experiments with ex vivo samples, on the other hand, do not suffer from this limitation, allowing long scanning sessions without data degradation caused by motion and for histology to be performed as a means of validation of any microstructural findings (20,21). One shortcoming of ex vivo experiments is that MR measured parameters in tissues undergone formalin fixation (a common practice to preserve human post-mortem tissue) show substantial differences to those found in vivo. Those differences have been seen both on single- and multi-compartment relaxometry, and diffusion-weighted imaging (22,23). Yet, a previous study showed that the bulk magnetic susceptibility of brain tissues measured by QSM did not change significantly between in vivo and ex vivo conditions, and also during a 6-week fixation period (24). This finding is in agreement with the experiment results when studying the microstructural effect in phase imaging (25), where comparable bulk magnetic susceptibilities were observed between fresh and fixed rat optic nerves but the origins of the susceptibility contrast are different.

In this study, we investigate the magnetic susceptibility, compartmentalisation and chemical shift effects on the MR phase using a formalin-fixed, post-mortem human brain specimen for WM phase-contrast mechanism studies at 3T, providing comprehensive insights for the use of fixed tissue in future QSM methodology studies. We performed multiple orientation experiments in both whole-brain and excised tissue samples (26). This enabled us to obtain both traditional QSM maps and ground truth bulk magnetic susceptibility measurements of the excised samples, as well as a separate estimation of microstructure compartmentalisation information. The samples were then studied using electron microscopy (EM) to further evaluate microstructural correlates between MR and histology.

## Methods

The study is divided into 3 parts:

1. a post-mortem, formalin-fixed human brain specimen was scanned on an MRI scanner in multiple orientations with respect to B_0_ for robust magnetic susceptibility measurements;
2. homogeneous WM specimens were excised from the whole-brain specimen, embedded in agar and subsequently scanned again in various orientations in respect of B_0_ to measure their magnetic susceptibility and microstructure-induced field, similar to the experiment conducted by (26);
3. the WM specimens used in the second part of the study were imaged by 3D EM, allowing MRI data to be compared to histology.

### Tissue processing

A post-mortem human brain specimen from a deceased male (aged 78 years old) with no history of neurological disorder (cause of death: myocardial infarction) was used for this research in accordance with the local ethics committee and the Anatomy Department of Radboud University Medical Center (Radboudumc, Nijmegen, the Netherlands). The brain specimen was immersed in 10% formalin for tissue fixation after being extracted from the skull.

After one month of fixation, the specimen was scanned on an MRI scanner (imaging details in section 2.2) to obtain the brain morphology for creating a tailor-made holder. The holder was made of a stack of 35 4-mm thick, 3D-printed plastic plates having space with the same shape of the specimen in the centre, covering most of the brain (see Figure 1) where the specimen can be fitted tightly inside the holder and a surrounding spherical container. Each plate has a grid layout with 4 mm x 4 mm elements, providing landmarks in MRI images for planning and guidance of tissue excision for the validation experiment in the second MRI session.

**Figure 1:**
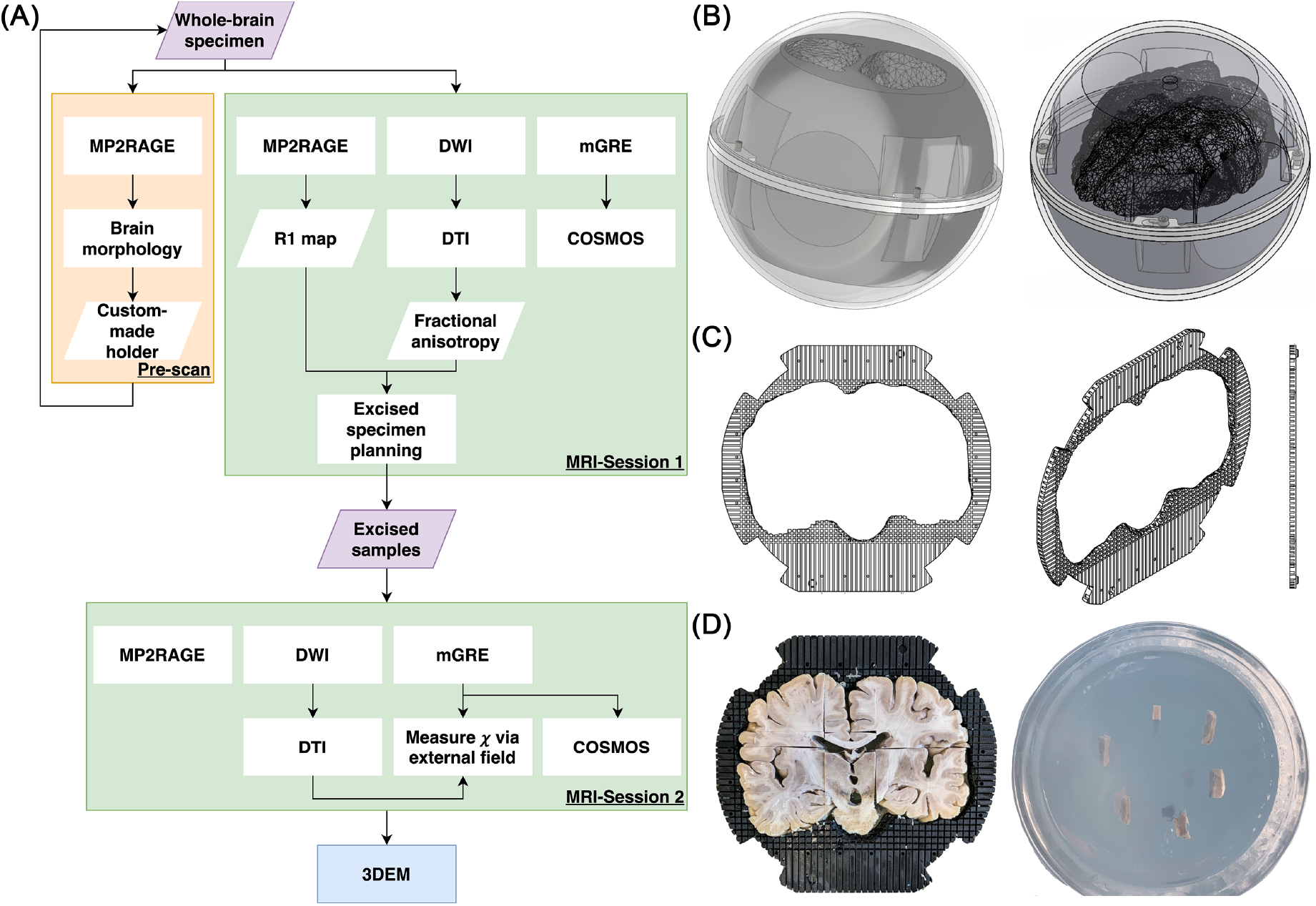
(A) A summary of this study, consisting of a pre-scan, two MRI sessions and a 3D EM session. (B) Schematic of the experimental setup used in the first MRI session. The setup is made of two parts: an outer transparent sphere allowing free rotation of the specimen and a tailor-made inner holder to ensure the specimen was in a fixed position inside the sphere. (C) Schematic of the plate which forms the inner holder. The centre of the plate is a space with the shape of the specimen, surrounded by a grid structure providing location reference in MRI image (the gaps filled with water while the plate material gave no detectable signal), and guidance in sample excision for the second imaging session. (D) An illustration of how homogenous samples were acquired with the aids of the plate (right) and the excised specimens were embedded in the agar inside the cylindrical container (left).

After the first MRI session, 10 homogeneous WM regions of interest (ROIs) were identified (DTI fractional anisotropy ≥ 0.45) with excisable volume (≥ 1 element of the holder plate grid, i.e., 64 mm^3^). These WM tissues and 2 additional deep GM tissues (one from globus pallidus and one from putamen) were then excised and embedded in 1% low-gelling temperature agarose (A9414, Sigma Aldrich, Germany; with demineralised water) to ensure no extra tissue protein denaturation would occur between the two acquisitions. The samples were positioned in a polymethyl methacrylate cylindrical container with the main fibre orientation perpendicular to the cylindrical axis. Imaging was performed about 1 day after the excised specimens were acquired and 5 days after the whole brain scan.

EM was utilised to provide an additional reference to understand and explain the MRI findings. Two days after the second MRI session, the WM specimens were sectioned to 100 *μ*m on a vibratome (VT1000S, Leica Biosystems, Nussloch, Germany) before being immersed in 2.5% glutaraldehyde in 0.1M sodium cacodylate buffer for overnight incubation at 4°C. The specimens were then transferred to 0.25% glutaraldehyde in 0.1M sodium cacodylate buffer for storage at 4°C, and then delivered to the EM facility at the University of Oxford for imaging. The workflow of this study is summarised in Figure 1.

### MRI experiments

#### Data acquisition

The study was approved by the local ethics committee. All MRI data were acquired on a 3T scanner (Prisma, Siemens, Erlangen, Germany) at room temperature (20°C) using a 64-channel array head/neck coil (with only 48 head channels were enabled). The experiment consisted of two imaging sessions: the first session was conducted on the whole-brain specimen and the second session was conducted on the excised brain tissues. The following protocol was used for the first session:

1. MP2RAGE adopted to sensitise for T_1_ values between 250 ms and 1000 ms, 1 mm isotropic resolution, TI1/TI2/TR=311/1600/3000 ms, flip angle (*α*) #1/#2 = 4°/6°, Tacq = 5 min;
2. 2D spin-echo EPI DWI, 1.6 mm isotropic, TR/TE=15241/77.6 ms, 2-shell (b=0/1250/2500 s/mm^2^, 17/120/120 measurements with 7 b=0 measurements collected with reversed phase-encode blips for distortion correction), 20 repetitions, Tacq = 5.6 hours;
3. Monopolar, 3D multi-echo GRE, 1 mm isotropic resolution, TR/TE1/ΔTE/TE6=40/3.45/6.27/34.8 ms, *α* =20° (optimised for WM T_1_), 10 orientations with respect to B_0_, chosen to optimise microstructural information decoding (27); Tacq = 1.7 hours.

In the second scanning session, the excised specimens were scanned with the following protocol:

1. MP2RAGE sequence with the same parameters as above;
2. 2D spin-echo EPI DWI, 1 mm isotropic, TR/TE=15241/77.6 ms, 2-shell (b=0/1250/2500 s/mm^2^, 17/120/120 measurements with 7 b=0 measurements), 9 repetitions, Tacq = 9 hours;
3. Monopolar, 3D multi-echo GRE, 0.7 mm isotropic resolution, TR/TE1/dTE/TE6=40/3.45/6.27/34.8 ms, FA=15° (optimised for agar T_1_), 10 orientations with respect to B_0_ (acquisition order was randomised); Tacq = 5.3 hours.

#### Data processing

Each DWI repetition was pre-processed separately with MP-PCA denoising (28), susceptibility-induced distortion correction (29,30) and eddy current-induced distortion correction (31). DTI model was then utilised on the DWI data after averaging all repetitions. R_1_ maps and DTI results were then linearly registered to the GRE data using ANTs (32).

GRE data with all orientations were first corrected for gradient nonlinearity induced geometric distortion and then linearly registered to a common space, independent of the experiment orientations. R_2_* maps were computed using a closed-form solution (33). Field maps were computed using SEGUE (34) spatial phase unwrapping with optimum-weighted echo combination (35) and tissue field maps were computed using LBV (36) in SEPIA (37). For the whole-brain data, bulk isotropic magnetic susceptibility was derived using COSMOS (38). Additionally, the QUASAR algorithm for multi-orientations was also applied to test if the bulk isotropic magnetic susceptibility measurement improved when the field generated by non-susceptibility contributions (f_*ρ*_) were simultaneously estimated (39):

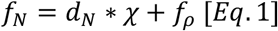

where f_N_ and d_N_ are the tissue field and a unit dipole field associated with the acquisition at orientation N and ∗ is the convolution operator.

For the excised specimen data, in addition to the computation of the bulk isotropic magnetic susceptibility using COSMOS, quantification of isotropic and anisotropic magnetic susceptibility (*χ*_i_ and *χ*_a_) of the sample without confounding with non-susceptibility microstructural contributions was performed by fitting the measured field f_N_ on the agar surrounding the specimen in the following manner (26):

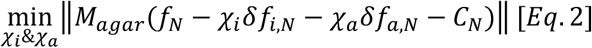

where: M_agar_ is the binary mask on agar with inner and outer boundaries 1 and 5 voxels away from the specimen tissue boundary in all directions; *δ*f_i,N_ and *δ*f_a,N_ are the frequency perturbations generated by a homogeneous specimen per unit of isotropic and anisotropic magnetic susceptibility at orientation N, with its orientation of tensor derived from obtained from DTI; and C_N_ accounts for any baseline frequency differences in agar due to either chemical exchange in the agar compartment or any residuals remaining after background field removal for a particular orientation. Linear regression analysis was then conducted to compare the magnetic susceptibility measurements between the COSMOS and external field method, and between the two imaging sessions to determine if the excised specimens agree with the ROIs in the whole-brain imaging data.

Tissue compartmentalisation contributions to the MR phase can be measured as the residual field (f_R_) inside the specimen with mask M_specimen_:

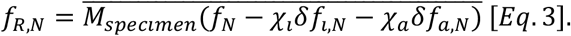

which is expected to vary with the angle *θ* between the fibre direction and B_0_ as

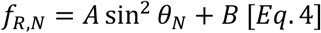

where A accounts for the microstructure orientation-dependent effect of the specimen and B is orientation-invariant, related to both magnetisation exchange and microstructure (26).

### Electron microscopy

Two corpus callosum (CC) specimens from the second MRI session (CC4 and CC5) having the greatest discrepancy of the microstructural compartmentalisation effect underwent 3D EM (40) to provide histology data for the MRI experiment validation. Each of the EM images has a matrix size of 8000×8000 with an in-plane resolution of 13.7×13.7 nm (~0.11×0.11mm FOV) and slice thickness of 100 nm. In total, 651 and 623 slices were acquired for CC4 and CC5 respectively.

Our EM data showed similar myelin sheath damage (splitting and swelling) as illustrated in (41), resulting in unsatisfactory compartmental classification (intra-axonal, myelin and extra-axonal compartments) from standard segmentation tools. To facilitate high-quality 3-compartment classification, we first performed a semi-automatic intra-axonal segmentation using ITK-snap (42) on down-sampled 3D EM data (87.7×87.7 nm and 100 consecutive slices). The myelin sheath of each axon was initially defined by expanding the axon assuming a g-ratio = 0.5 (for axon diameters<1.2*μ*m) or 0.6 (otherwise), followed by intensity thresholding using the EM images. The resulting myelin mask is clearly influenced by the chosen threshold, therefore, myelin volume fractions (MVF) derived from 5 threshold values range from 125 to 145 (step size of 5, most frequent intensities in the myelin mask are 104 (CC4) and 88 (CC5)) were reported. To account for the enlarged myelin volume due to swelling, we also computed the image intensity corrected myelin probability as:

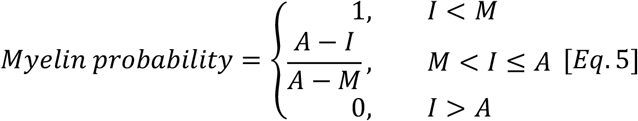

where A and M are the most frequent values inside the axonal and myelin masks and I is the intensity of a voxel. The axon and myelin masks of each myelinated axon were eventually combined in a single classification map to avoid repeated counts due to overlapping between neighbouring axons.

Axonal volume fraction (AVF) and MVF were computed by counting the total number of voxels of each compartment in the classification map, and were used to derive the sample g-ratio (20). Effective axonal diameter was defined as the square root of the product of the 2^nd^ and 3^rd^ principal axis lengths of the axons obtained from the regionprop3 function of MATLAB (Mathworks, Natick, US), from which the median and the skewness of the axonal diameter distribution were computed. Fibre dispersion was computed from the axonal volume-weighted average squared dot product between the axon main principal direction and the average orientation of the entire sample (27). To further investigate the effect of realistic myelin sheath geometry on the compartmental frequency shifts, field perturbations induced from myelin ***χ***_i_ was simulated in two scenarios: when the segmented sample axons were parallel or perpendicular to B_0_. The frequency shift distribution in the extracellular space was subsequently analysed.

## Results

Whole-brain imaging results are shown in Figure 2. The R_1_ maps obtained from the first session (5-month fixation) show faster relaxation rates than those from the pre-scan (1-month fixation), with the contrast between WM and cortical GM being clearly reduced, and DGM showing increased R_1_ (Figure 2A and 2B). The COSMOS derived magnetic susceptibilities are in reasonable agreement with previously published in vivo data (see supplementary Figure S1), where opposite magnetic susceptibility between WM and GM can be observed (Figure 2C). However, the residual field of the COSMOS estimation shows a slowly-varying pattern across the brain that cannot be explained by the isotropic dipole field (Figure 2E) and is relatively stable across orientations (see Figure 2F). This residual map shares similar contrast and values with the QUASAR non-susceptibility contributions map (Figure 2H). Hence, not surprisingly, the bulk magnetic susceptibilities derived from QUASAR (Figure 2G) and COSMOS (Figure 2c) are comparable. Susceptibility tensor imaging was also performed, yet the results beyond the mean susceptibility tensor were found not informative.

**Figure 2:**
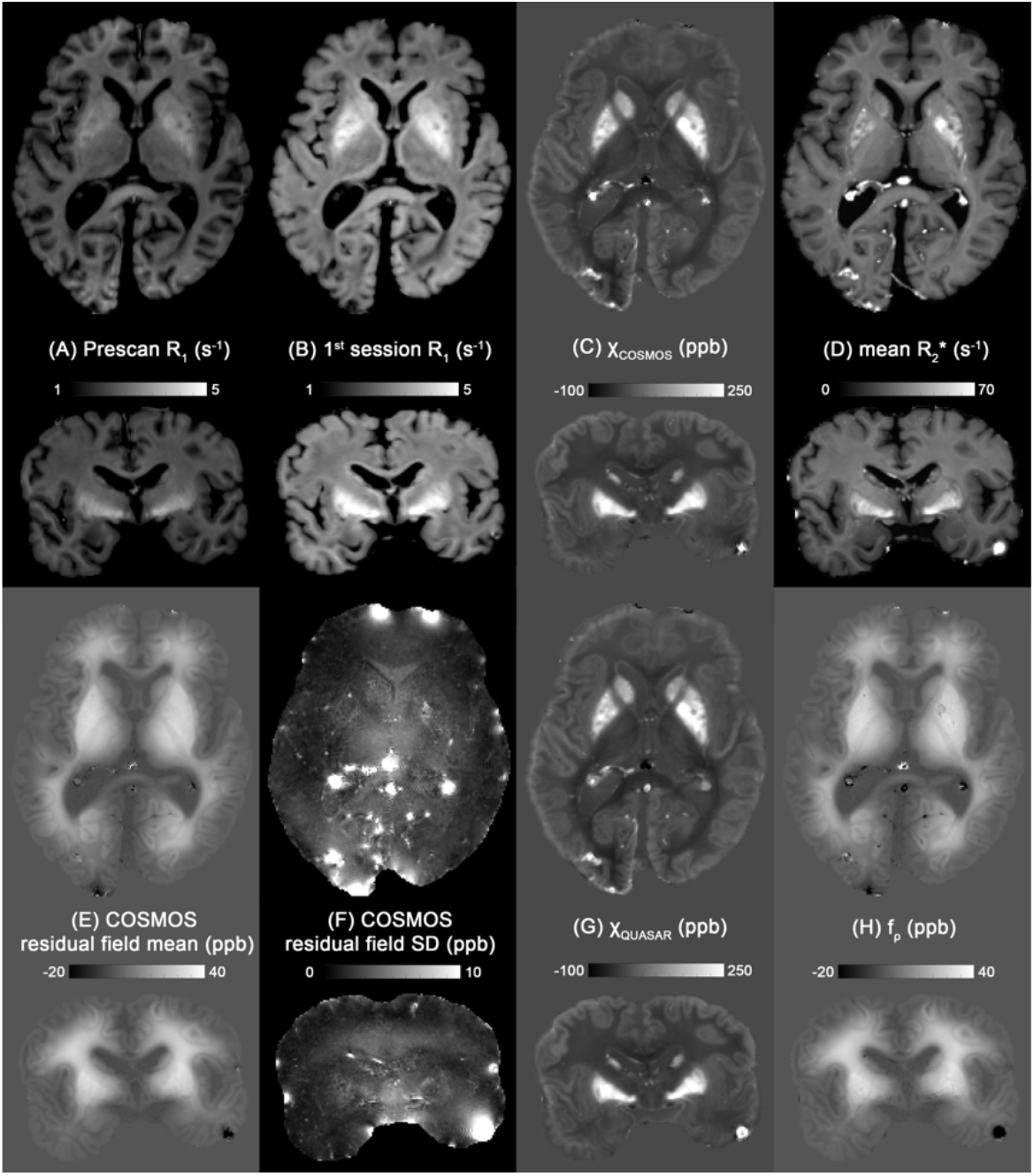
Quantitative maps of the whole-brain specimens in transverse and coronal directions. (A) R_1_ map obtained from a pre-scan after 1-month of formalin fixation and (B) R_1_ map obtained from the 1^st^ imaging session while the brain was fixed for 5 months. (C) COSMOS derived bulk magnetic susceptibility, (D) mean R_2_* map across the 10 rotations, (E and F) mean and standard deviation of the residual fields from COSMOS across orientations, (G and H) QUASAR derived bulk magnetic susceptibility map and non-susceptibility contribution map.

Figure 3 shows the magnetic susceptibility of the excised specimens measured via the external field on agar with their ROIs illustrated in the whole-brain R1 map. The mean *χ*_i_ and *χ*_a_ are −1.17±9.18 ppb and 4.03±1.63 ppb across WM specimens. A relatively strong positive *χ*_i_ is found in the corticospinal tract specimen (CST; 19.17 ppb), which was found in retrospect to be due to some DGM residual in one end of the excised sample. The coefficient A of sin^2^*θ* dependence reflecting the WM microstructure effect has a mean of 1.46±1.55 ppb with a mean intercept B of −2.75±0.79 ppb in WM. However, the R^2^ of the specimen residual field fitting suggests that not all WM specimens fit the sin^2^*θ* function equally well, particularly for samples obtained from the genu and splenium of the CC (CC7-CC9; R^2^ ranging from 0.01 to 0.71). Therefore, we focused on the 6 WM specimens obtained from the body of the CC (CC1-CC6) in comparison to the whole-brain data in the linear regression analysis.

**Figure 3:**
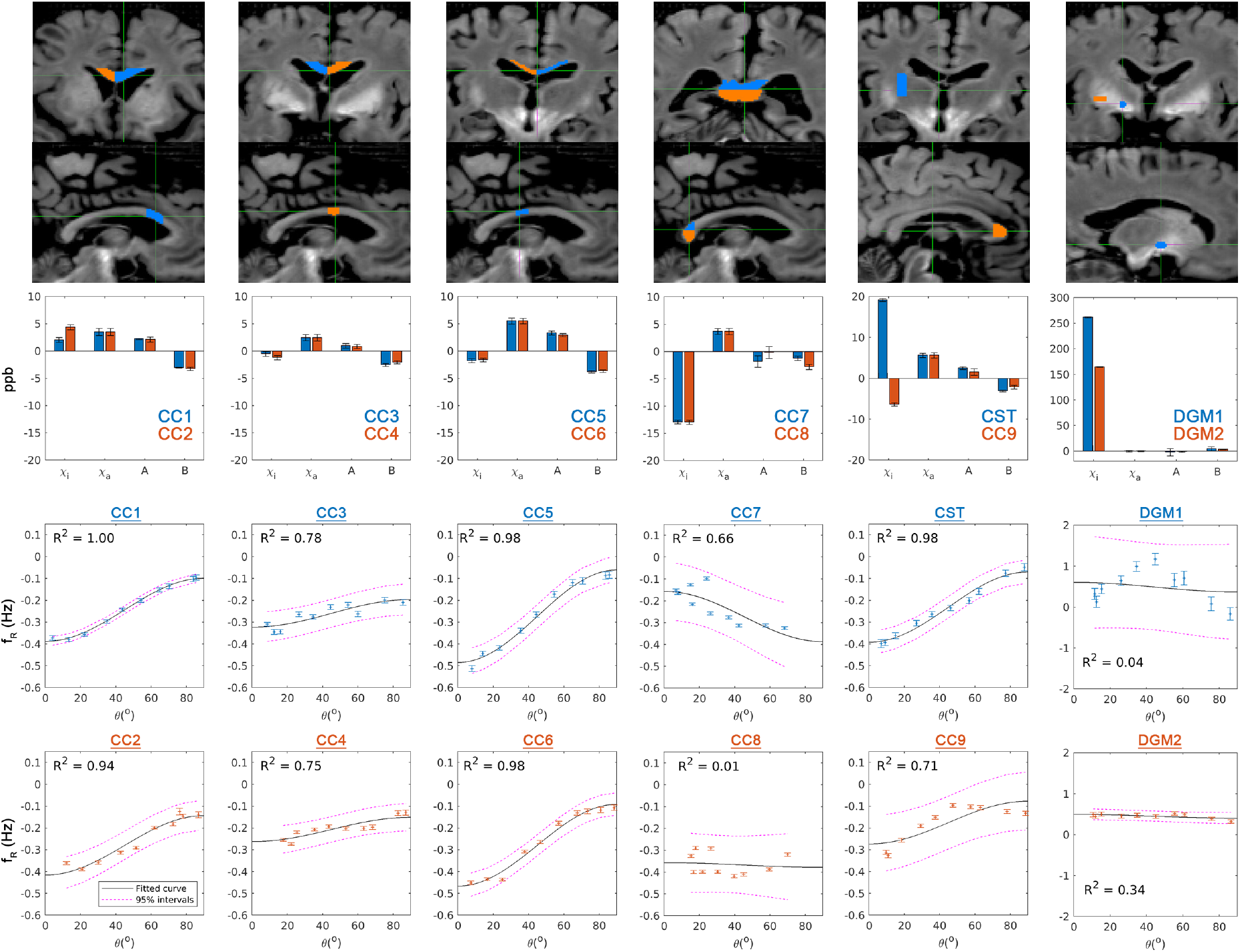
(Middle row) Barplots of isotropic and anisotropic magnetic susceptibility, and coefficients A and B of the fitting of sin^2^*θ*. (Bottom two rows) Fittings of sin^2^*θ* reflecting microstructure compartmentalisation of the excised specimens. Each column shows the results of two specimens with their ROIs illustrated in the whole-brain R_1_ map (top two rows). Error bar indicates the standard error (except for the coefficients A and B in the barplots; 95% confidence intervals in this case). CC: corpus callosum; CST: corticospinal tract; DGM1: globus pallidus; DGM2: putamen.

Strong linear relations were found in mean susceptibility estimated by COSMOS between the excised specimens and the corresponding ROI in the whole-brain data (cross-session; R^2^=0.603, Figure 4A), between the *χ*_i_ from external field measurement and the mean COSMOS susceptibility in the whole-brain data (cross-session, cross-method; R^2^=0.783, Figure 4B), and between the *χ*_i_ from external field measurement and the mean COSMOS susceptibility on the excised specimens (cross-method; R^2^=0.925, Figure 4C). All the slopes of the linear regressions are close to 1, whereas the relatively large intercepts in Figure 4A-B reflect the different reference medium in the scans (1^st^ session: water; 2^nd^ session: 1% agar).

**Figure 4:**
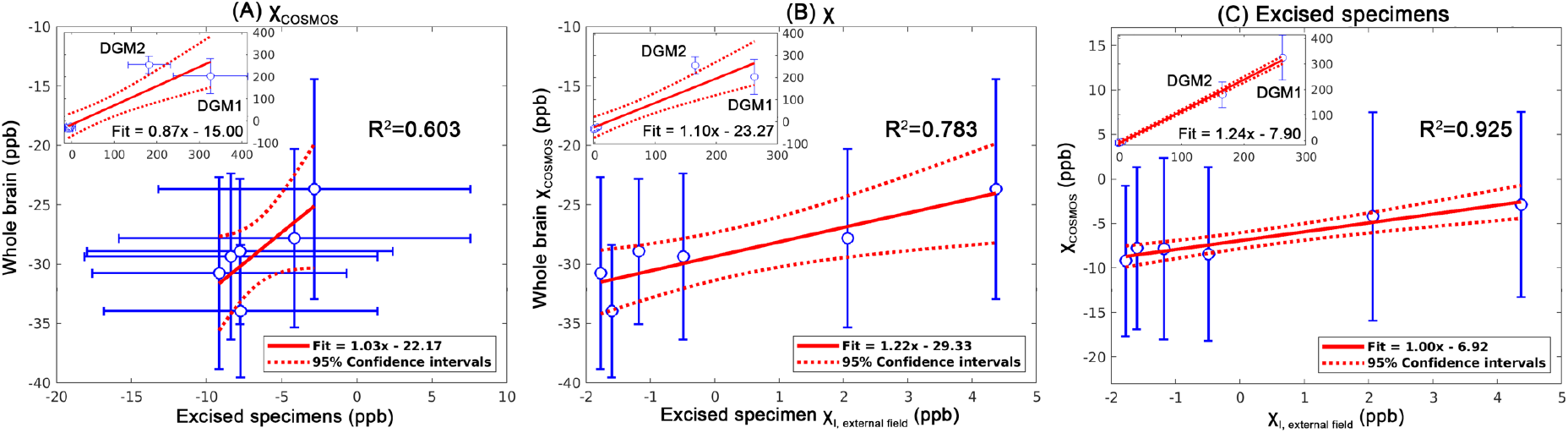
Linear regression analyses on the 6 WM specimens of (A) the magnetic susceptibility measured between two imaging sessions using COSMOS, (B) between the bulk magnetic susceptibility measured by COSMOS on the first session and the *χ*_i_ from external field measurement on the second session, and (C) the excised specimens between COSMOS magnetic susceptibility and external field derived *χ*_i_. Blue points: measurement data; solid red line: fitted line; dash line: 95% confidence interval; error bar: standard deviation. The subplot of each panel shows the regression result when the DGM specimens were included.

The microstructural properties of two WM specimens (CC4: relatively weaker microstructural phase; CC5: strong microstructural phase) derived from 3D EM data are summarised in Figure 5. Both specimens have similar MVF (8% difference), AVF (4% difference) and axonal diameter (1% difference). Noticeable differences are observed in fibre dispersion (50% difference) and in the FWHM of the extra-axonal frequency distributions when the fibre direction is parallel to B_0_ (63% difference), the latter being attributed to the differences of both fibre dispersion and the spatial distribution of myelinated axons.

**Figure 5:**
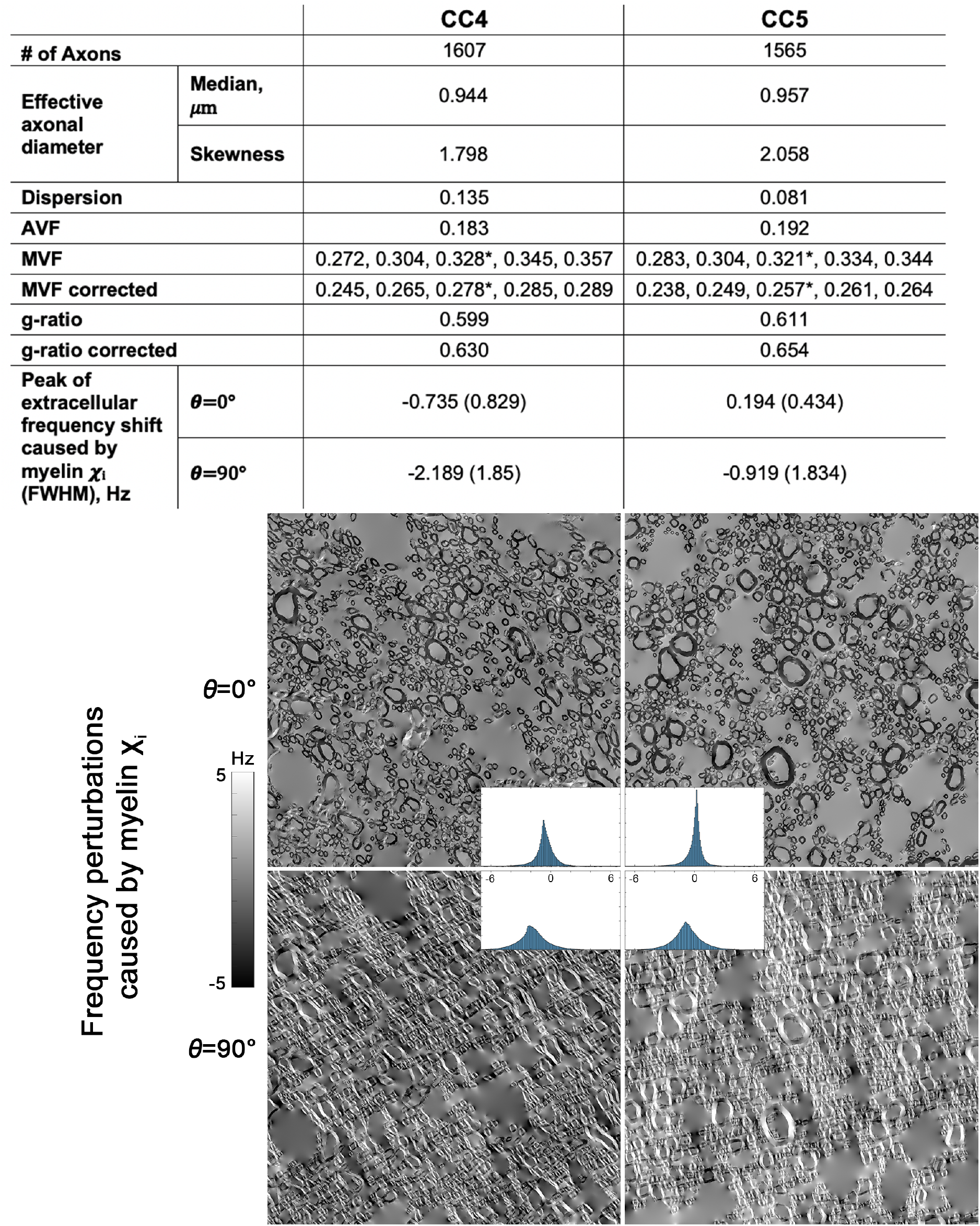
(Top) Summary of the 3D EM derived CC4 and CC5 microstructural properties. MVF was separately probed using 5 different intensity thresholds (only the middle value indicated by * was showed in g-ratio). The corrected MVF was derived using Eq. [5]. (Bottom) Frequency induced by the myelin ***χ***_i_ at two orientations to B_0_ (***θ***=0° & 90°). Sub-figures show the frequency distributions in extracellular space, where the locations and the FWHM of the peaks are shown in the above summary.

## Discussion

In this study, we examined the magnetic susceptibility and microstructural compartmentalisation effect on MRI phase data on a formalin-fixed, post-mortem human brain specimen. The bulk magnetic susceptibility of the whole-brain specimen shows comparable contrast to those in the previous ex vivo studies (6,24), as well as to in vivo imaging: WM is slightly diamagnetic, whereas cortical and deep GM are paramagnetic. Further investigation reveals the residual fields of COSMOS have a gradient-like appearance varying from the surface toward the centre of the specimen (Figure 2E), which is similar to the expected way of how the solutions (fixative or water) diffused into the specimen. Since these residual fields are relatively constant across different rotations, it is likely to be caused by the exchange effect (17–19). These fields were captured as the non-susceptibility contributions by QUASAR and they did not have a significant impact on the bulk magnetic susceptibility measurement (Figure 2C, 2G). This result is distinct from in vivo imaging results (Figure S1), where the susceptibility differences in WM are more noticeable, suggesting that the effect of (sub)cellular structure of WM is considerably reduced and the sphere of Lorentz inclusion utilised in COSMOS is already a good approximation on formalin-fixed tissue.

While the bulk susceptibility of the WM samples in this study is similar to in vivo imaging, the residual field analysis inside the excised homogenous tissue confirmed that the microstructure compartmental frequency (parameter A in Eq. 4) is notably weaker in our samples than in vivo and reported by others. In a similar experiment (26), the amplitude of the microstructure frequency of a fresh bovine optic nerve at 7T was −18.75 ppb, significantly larger in magnitude and with an opposite sign to what we have obtained in our CC samples, 1.46 ppb. Additional analysis was performed to consolidate this result (see supplementary Figure S2). A reduction of the microstructural compartmentalisation effect had already been reported in the literature when studying fresh vs fixed rat optic nerves (25). One possible explanation is the structural alteration of the myelin sheath in fixed tissues. In our 3D EM images, we observed myelin sheath spitting and swelling in some of the myelinated axons, similar to the observation reported in the previous study (41), and such phenomena appeared more frequently in larger axons than small axons. Based on the general Lorentzian tensor approach (43), the increase of the aqueous space of the myelin sheath can result in the amplitude reduction of the induced frequency shifts inside the myelin sheath and the intra-axonal space. Microstructural differences related to structures (bovine optic nerve vs human CC) and age-associated demyelination (45,46), together with the tissue preparation methods can also contribute to the differences observed in this study, as all these factors modulate the relative water concentration in the three WM compartments.

All six specimens obtained from the body of the CC have similar magnetic susceptibility anisotropy, suggesting that they have similar MVF based on the HCM approximation (Eq. S25 in (17)), and this is supported by the EM analysis (0.278 & 0.257 between the two samples). The amplitude of the residual field inside the specimens is, on the other hand, subject to various properties including MVF, AVF and the aggregate g-ratio of the sample (Eq. A14 in (16)). Interestingly, the realistic geometry of the WM fibre also plays an important role in the compartmental frequency shifts (27,44). This effect is clearly illustrated in the frequency perturbation simulations in Figure 5: not only the centres but also the FWHM of the extra-cellular frequency distribution of the two samples are different, despite the two specimens having virtually identical MVF and AVF. The broader frequency spectrum of CC4 induces a faster R_2_* decay in the extra-axonal space and the specimen also has a more disperse fibre arrangement. These two factors together reduce the amplitude discrepancy between the slow R_2_* (intra- and extra-axonal water) and the fast R_2_* (myelin water) compartments throughout the echo time as well as the frequency difference between those compartments that could result in a reduced compartmentalization effect.

The experimental setup, particularly the 3D-printed holder, is an effective tool also for other ex vivo studies that involve histology. The grid on the 3D-printed plates not only facilitates tissue excision with high precision but also provides landmarks in MRI images for experiment planning and sample matching. The close to unity linear relations of the susceptibility measurements between the two sessions (Figure 4A and 4B) support that the ROIs drawn on the whole-brain images and the excised specimens are corresponding to each other.

One limitation of this study is the relatively long fixation time of the specimen. The first imaging session happened after 5 months of fixation instead of the scheduled 2 months because of the coronavirus measures that took place in the Netherlands. The prolonged fixation time results in a substantial change of specimen R_1_ between the pre-scan and the first session (Figure 2A, 2B; average R_1_ across the brain increased from 1.95±0.47 s^−1^ to 2.67±0.74 s^−1^). The enhanced R_1_ in deep GM suggests the contribution of iron to the R_1_ was more pronounced. An alternative and more likely explanation would be that the myelin contribution to R_1_ could be reduced by the fixation process, which would also explain the diminished cortical GM and WM contrast observed at 5 months. Experiments conducted with a shorter fixation time can potentially reduce the fixation effects on the signal phase. However, progressive changes in relaxation parameters begin in the early stage of fixation (23) and it is likely that fixation induced phase differences could also happen simultaneously. Lastly, the EM analysis can be subject to sampling bias due to the limited field-of-view. Preliminary results of a recent study suggest that large axons in the WM tissue can be under-represented in EM compared to light microscopy when larger field of view is available (45). Such under-representation may have an impact on the realistic geometry of the myelinated axons on the phase data.

## Conclusion

The contributions of MR phase contrast observed in the formalin-foxed brain specimen are substantially different from fresh tissue, despite the QSM maps derived from in vivo and ex vivo imaging sharing similar contrasts and values. Particularly, the reductions of magnetic susceptibility anisotropy and compartmentalisation are observed in the fixed WM tissue. An increase of non-susceptibility contributions to phase contrast can also be found in fixed tissue, which is potentially introduced by formalin fixation. Therefore, WM magnetic susceptibility and microstructural quantification findings in studies using formalin-fixed tissue should be interpreted with care. Our study suggests that the microstructural effects observed in our samples encode information regarding WM arrangements such as dispersion and packing while susceptibility anisotropy encodes myelin volume as was predicted from theory.

## Acknowledgement

This work was funded by the Netherlands Organisation for Scientific Research (NWO) with project number FOM-N-31/16PR1056. The authors would like to thank Ms. Sibrecht Bouwstra for the 3D-printed holder construction, Ms. Shaghoyegh Abghari for laboratory support, Dr. Errin Johnson for 3D EM data acquisition, and Dr. Ferdinand Schweser for the fruitful discussion on magnetic susceptibility analysis including chemical exchange terms.

## Notes

### Competing Interest Statement

The authors have declared no competing interest.

