## Supplementary material for "Studying magnetic susceptibility, microstructural compartmentalisation and chemical exchange in a formalin-fixed ex vivo human brain specimen"

### Applying COSMOS and QUASAR on in vivo imaging dataset

Figure S1: COSMOS and QUASAR results on (left) in vivo imaging dataset from QSM challenge 1 and (right) formalin-fixed post-mortem brain specimen. (From top to bottom) Bulk magnetic susceptibility (𝝌) maps derived by COSMOS, 𝝌 maps derived from QUASAR, differences between the COSMOS and QUASAR 𝝌 maps, and non-susceptibility contribution maps derived from QUASAR. Comparable imaging contrasts can be observed between the in vivo and ex vivo imaging results, where iron-rich basal ganglia such as globus pallidus, red nuclus and substantia nigra are the brightest in the maps (blue arrows) whereas myelin-rich white matter is dark. Interestingly, the in vivo QUASAR derived 𝝌 map is more homogenous within white matter in contrast to the COSMOS counterpart (red arrows). The contrast among WM fibre bundles in the 𝝌 map can be originated from magnetic susceptibility anisotropy and microstructural difference and is partially explained in the non-susceptibility contribution map of QUASAR. The difference in 𝝌 between COSMOS and QUASAR is considerably smaller on our formalin-fixed specimen compared to those derived from the in vivo dataset.

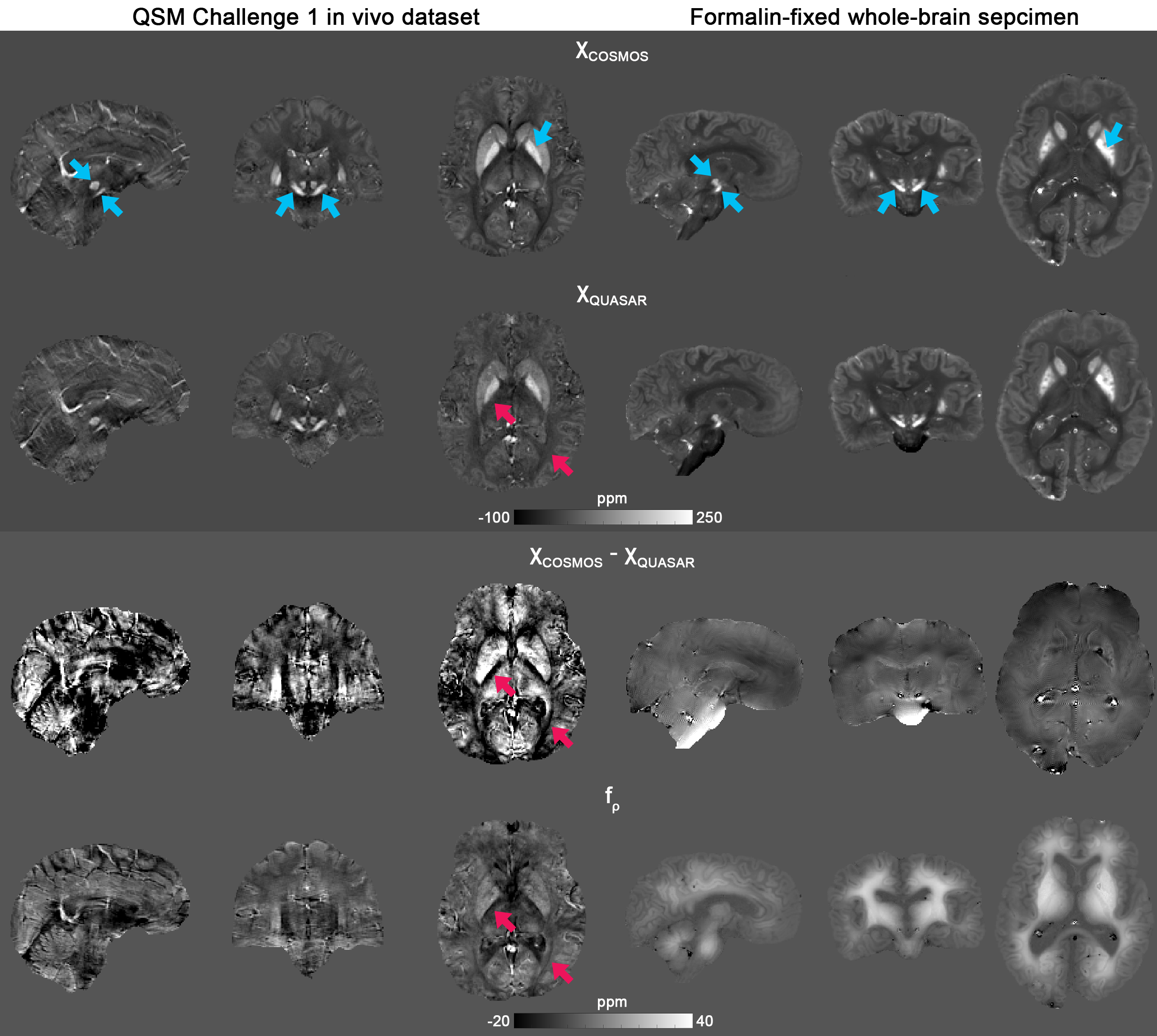

### Measurement of isotropic and anisotropic magnetic susceptibility based on external field of the homogenous white matter samples

Figure S2: Results of fitting the isotropic and anisotropic magnetic susceptibility of the homogenous, excised WM tissue specimens (top 2 rows) with and (bottom 2 row) without a constant term to account for acquisition difference (e.g. shimming) for each orientation. Blue line represents the mean residual field in the external agar region included to compute the susceptibilities; red line represents the mean measured local field; and yellow line represents the fitted constant terms. Note that the yellow lines have an identical shape as the mean residual field when the constant term was not included in the fitting, and the mean residual fields are close to zero once we introduced this term in the fitting.

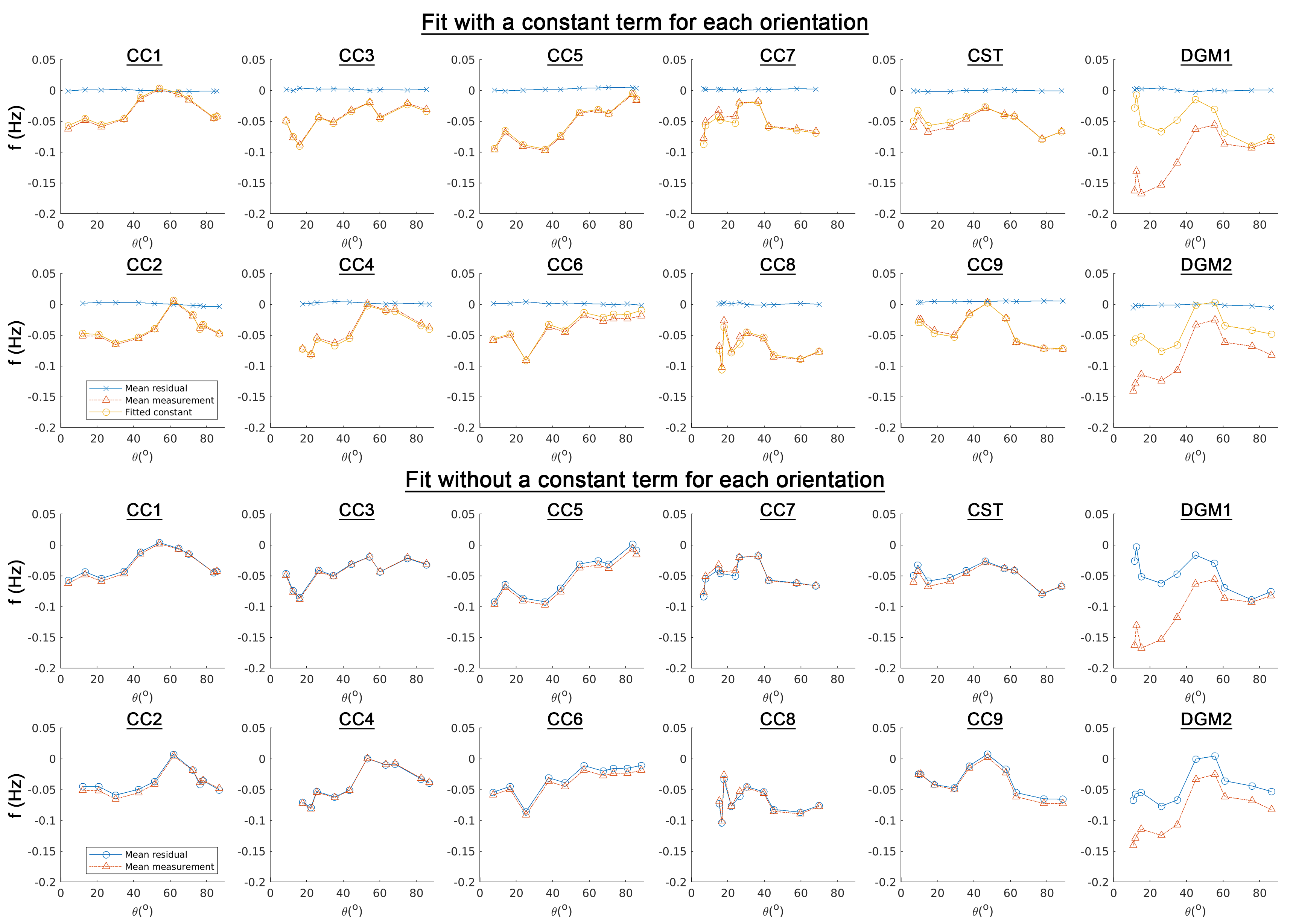
